# SPARQ-MI leverages end-to-end spatial single-cell analysis of the tumor microenvironment

**DOI:** 10.64898/2026.06.06.730569

**Authors:** Lukas Kiwitz, Roberta Turiello, Maike Effern, Marieta Toma, Jennifer Landsberg, Michael Hölzel, Kevin Thurley

## Abstract

Detailed spatial analysis of the tumor micro-environment (TME) through multiplexed fluorescence imaging requires quantitative image-processing and data-analysis methods. While data-preprocessing down to segmentation of individual cells is captured by available methods, statistical analysis of single-cell features is compromised by the uneven noise distribution especially in complex tissues such as the TME, as well as by labor-intensive manual cell-type annotation and region segmentation. Here, we present SPARQ-MI (Spatial Phenotyping, Architecture Reconstruction and Quantification from Multiplexed Imaging) for streamlined spatial single-cell analysis, along with a tissue microarray PhenoCycler data-set with 37 fluorescent channels from melanoma patients under immunotherapy. We demonstrate that SPARQ-MI enables robust reconstruction of the cellular and spatial composition in this and other tissue types. Our analysis reveals associations of the cell-state and spatial location of CD8 T cells with response to immunotherapy. Overall, SPARQ-MI allows for quantitative analysis of complex fluorescence histology samples under minimal user input, and accounting for spatially uneven coverage of antibody signals, setting the stage for quantitative analysis of clinical samples.

## Introduction

The tumor micro-environment (TME) is a complex biological niche comprising tumor, immune and stromal cell types. Single-cell sequencing and immunohistochemistry have revealed connections between the composition and spatial organization of the immune-cell compartment and clinical outcomes in prevalent tumor entities, such as cholangiocarcinoma, renal cell carcinoma and melanoma. While cytotoxic CD8 T cells represent the dominant effector cell type in the TME, also helper T cells, regulatory T cells and various macrophage populations have been implicated in the anti-tumor response. Further, a robust anti-tumor immune response depends not only on effector cell abundance, but also on the location of immune cells within the TME(Lopez de Rodas *et al*, 2025; Mempel *et al*, 2024; van Weverwijk & de Visser, 2023). As a result, treatment efficacy for therapies that aim to reinvigorate the anti-tumor response has been linked to the cellular composition of the TME(Aliazis *et al*, 2025; Liu *et al*, 2023; Wang *et al*, 2023), and quantifiable TME states may serve as prognostic and predictive biomarkers enabling individualized treatment plans as well as advances in the mechanistic understanding of disease progression and new treatment options in solid tumors(Walsh & Quail, 2023; Engblom & Lundeberg, 2025; Elhanani *et al*, 2023). Comprehensive spatial profiling of the TME entails accurate detection and classification of specific immune-cell types in a dense and complex tissue context. Advances in multiplexed fluorescence imaging have made high-dimensional spatial proteomics datasets available for clinical research(Goltsev *et al*, 2018; Lin *et al*, 2016; Saka *et al*, 2019; Schubert *et al*, 2006; Wagner *et al*, 2019). Cyclic methods such as multiepitope ligand cartography(Schubert *et al*, 2006) (MELC), Co-detection by indexing(Goltsev *et al*, 2018) (Phenocycler) and Cyclic Immunofluorescence(Lin *et al*, 2016) (CycIF) repeatedly apply a low-plex staining protocol to the same tissue slide, while one-shot approaches such as PICASSO(Seo *et al*, 2022) capture a single multiplex panel and computationally separate fluorescence signals.

Leveraging the full capabilities of that technology for quantitative investigation of the TME requires robust image-processing and data-analysis approaches. Previously developed computational pipelines(Goltsev *et al*, 2018; Czech *et al*, 2019; Lu *et al*, 2022; Schapiro *et al*, 2022; Tan *et al*, 2025) focused on image pre-processing down to deep-learning based cell-segmentation models, which made imaging data accessible to single-cell analysis at scale(Bannon *et al*, 2021; Stringer *et al*, 2021). Nevertheless, a number of complications still compromise the accurate and efficient analysis of multiplexed microscopy data. Downstream of employing cell-segmentation models, single-cell mean-fluorescence intensity (MFI) is widely used to quantify staining intensity per cell and serves as primary feature used for cell-type annotation and characterization as well as spatial profiling. However, MFI is susceptible to batch effects between experiments such as spatial variation in staining intensity. Further, while in transcriptomics data analysis, cell-type annotation is often carried out through a combination of unsupervised clustering and manual annotation, multiplexed imaging-data requires strong over-clustering(Hickey *et al*, 2021), often making manual annotation a substantial bottleneck for the whole analysis workflow. Finally, unlike in non-spatial methods, detected cell counts depend on the specific sample micro-anatomy and are not comparable to measurements from homogenized tissues or between different tissue sections, and therefore specifically tailored approaches for tissue segmentation are required for systematic statistical analysis.

Here, we present SPARQ-MI (Spatial Phenotyping, Architecture Reconstruction and Quantification from Multiplexed Imaging) for quantitative analysis unifying image pre-processing, feature extraction, single-cell analysis and spatial analysis, along with a high-content Phenocycler data-set from 26 melanoma patients undergoing ICB therapy. SPARQ-MI provides natively normalized single-cell features, accurately detects antibody signal at single-cell resolution, and accounts for the sample micro-anatomy for patient-level analysis. An image-level feature-normalization approach achieves robust unsupervised classification of antibody-positive cells, and the algorithmic annotation of single-cell clusters enables deep phenotypic annotation. Employing SPARQ-MI for analysis of melanoma patients, we found that tumor-infiltration and the activation and cell-cycle state of specific immune-cell populations show strong predictive association with treatment response and patient survival.

## Results

### Quantitative analysis of multiplexed, high-content TME data

Multiplexed fluorescence imaging characterizes the complex cellular composition of immune-cell compartments *in situ*. Quantitative analysis is challenged by the large scale of input datasets as well as the inherent micro-anatomical variability of two-dimensional tissue sections, especially in the context of the TME. Here, we designed SPARQ-MI for image processing, single-cell annotation as well as spatial feature analysis and predictive association with clinical outcomes (**Figure 1A**). In order to process the large amounts of raw image data, we constructed a modular, parallelized, GPU-accelerated end-to-end pipeline that combines existing software libraries and tools with in-house solutions (**Figure 1B**). As input, the pipeline requires raw images and metadata, and as output, stitched multi-channel images, nuclear and cell-segmentation masks, and single-cell feature tables are generated (**Figure 1C**). The image pre-processing steps are based on the original analysis workflow for PhenoCycler data(Goltsev *et al*, 2018) and previously published pipelines and packages(Czech *et al*, 2019; Schapiro *et al*, 2022; Muhlich *et al*, 2022) such as Cytokit, MCMICRO, Ashlar, for image deconvolution, extended depth-of-field projection, tile stitching, cycle alignment and autofluorescence correction. Cell-segmentation is accomplished either using the Mesmer(Greenwald *et al*, 2022) model or by custom Cellpose(Pachitariu & Stringer, 2022) models. We calculate a network of physical cell-cell contacts based on the shared pixels in the segmentation mask. After image processing, cell segmentation and single-cell clustering, our method accurately detected antibody signals as well as spatial tissue architecture (**Figure 1D and E**) in murine spleen and clinical tumor samples.

**Figure 1:**
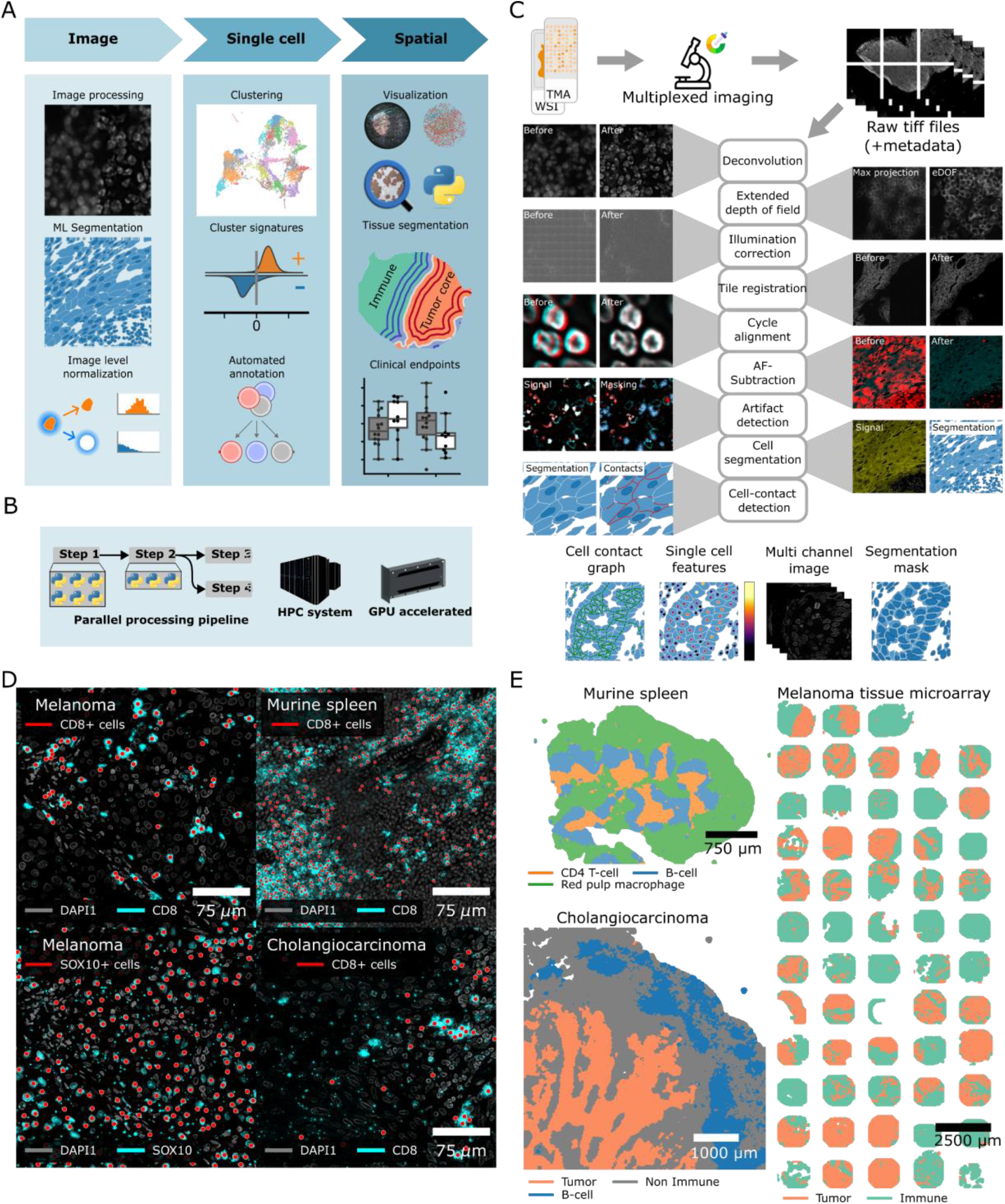
Overview of the multiplex image analysis pipeline. **(A)** Raw images are pre-processed, followed by application of deep-learning based cell-segmentation models. Additional steps for downstream analysis include image-level feature normalization and automated cell-type annotation. Spatial analysis is carried out by unsupervised tissue-segmentation, and sample-level features are used for predictive clinical modeling. **(B)** Schematic highlighting that our pipeline is modular, parallelized, and GPU-accelerated. **(C)** Schematic of the work-flow illustrating the effect of individual steps with representative images. The pipeline can take either raw whole-slide images (WSI) or tissue micro arrays (TMA) from CODEX devices as input data. **(D-E)** Example runs of murine and clinical tumor samples. Shown are (D) antibody-positive cells for selected channels, and (E) tissue annotation by dominant cell types.

### Feature normalization and single-cell pre-processing

A central problem in the analysis of fluorescence-microscopy based single-cell analysis is the fact that the fluorescence intensity of a cell may depend on its environment. Particularly dense cell-aggregates as well as off-target staining can bias a cell to appear brighter than an isolated cell (**Figure 2A**). Consequently, cell-type annotation and further cell-based analysis can be compromised compared to other sources of single-cell data such as flow-cytometry or transcriptomics, even if cell-segmentation is achieved with high quality. In order to address these challenges, we developed an image-level normalization scheme that adjusts MFI to the noise level in the local environment of each cell **(Figure 2B**). In a first step, we split the pre-processed images into foreground and background signal components using difference-of-Gaussian (DoG) filters. The filters are calibrated such that the background image approximates the noise distribution in the image, whereas the foreground image captures sub-cellular scale features. In a second step, we sample distinct foreground (*F*) and background (*B*) distributions for each cell to calculate the log2 fold-change (LFC) as

**Figure 2:**
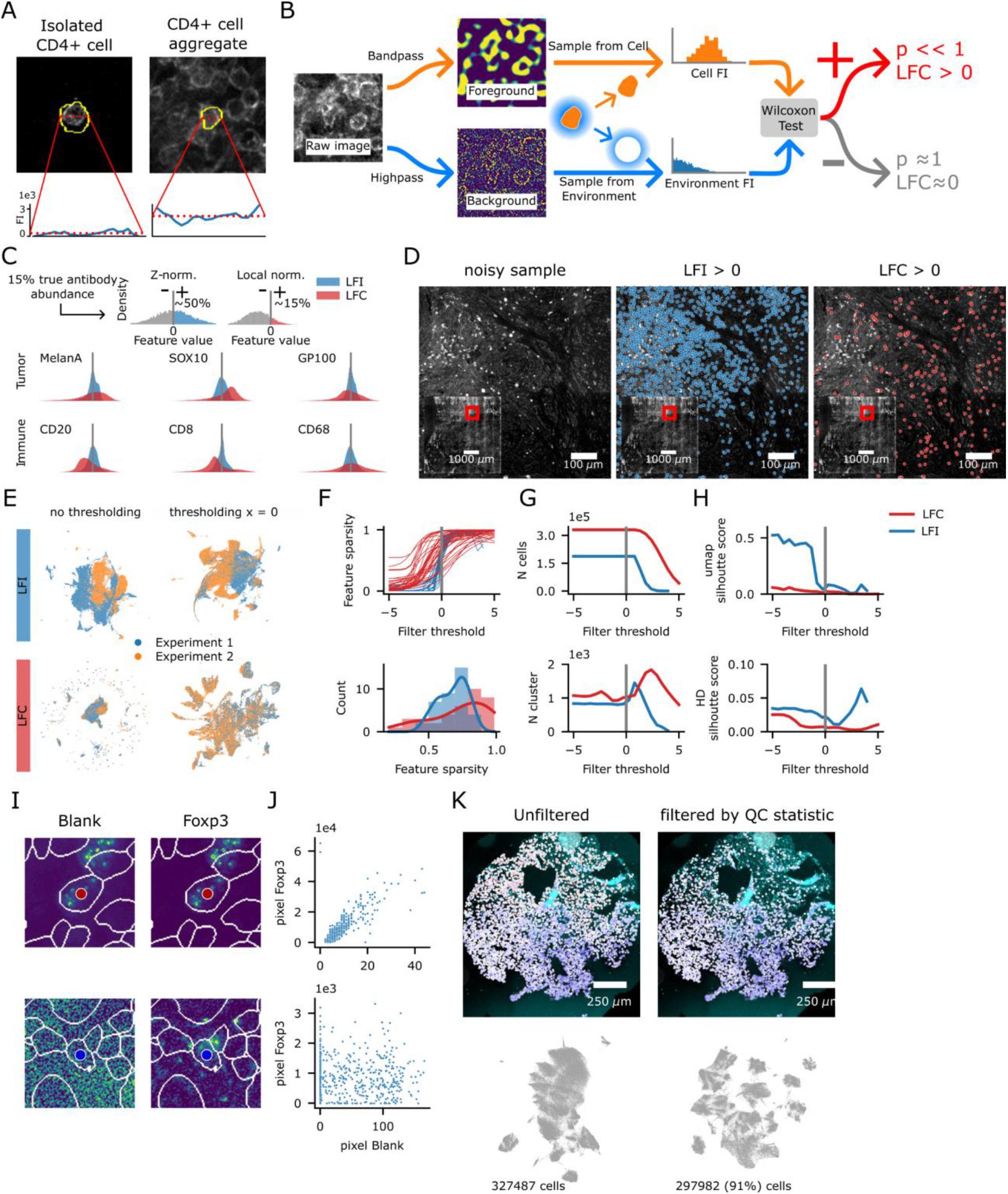
Image-level feature normalization. **(A)** Representative images from the melanoma TMA data-set indicating that MFI depends strongly on tissue location, as densely packed cells appear brighter than isolated cells of the same cell type. **(B)** Schematic of the feature-normalization workflow. Images are split into foreground and background frequency components using difference-of-Gaussian filters, and the log-ratio of local foreground and background signal (LFC) is exported as a single-cell feature. **(C-D)** Examples of (C) signal distributions and (D) image analysis, comparing the LFC feature to the conventional LFI feature, that is z-normalized MFI. The log LFC more accurately rejects false positive detections that arise as the consequence of structured noise patterns. **(E-H)** Comparison of downstream processing steps and validation statistics upon either LFC or LFI normalization, (E) thresholding of the feature matrix, (F) feature-vector sparsity and histogram of feature sparsity across antibodies for filter threshold x = 0, (G) fraction of cells retained after pre-processing and number of Leiden clusters with increasing filter stringency, (H) Measurements of batch effect in low-dimensional and high-dimensional (HD) space using cluster silhouette score. **(I)** Representative examples of cells with high (top) and low (bottom) correlation between antibody signal and blank channel, intensity ranges were scaled separately. **(J)** Association between blank and FoxP3 pixel intensities for the high and low correlation cell depicted in panel I. **(K)** Representative images (top) and umap embeddings (bottom) before and after selective filtering of autofluorescence-contaminated cells based on Spearman-correlation quality control statistics.

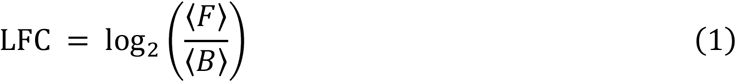

where ⟨*F*⟩ and ⟨*B*⟩ denote cell-wise average. The resulting LFC feature is inherently calibrated, where LFC = 0 indicates an equally distributed signal for a cell and its local environment. Further, based on the LFC, a one-sided p-value can be derived (**Figure S1A**), which is interpreted as a noise measure of the overall measurement and is useful for filtering out low-quality observations.

To assess the capacity of the LFC feature to represent biological information, we chose z-normalized logarithmic MFI (LFI) for comparison, as previously proposed(Hickey *et al*, 2021). For most antibody signals, the LFI distributions were centered around zero and appeared approximately unimodal, so that identifying cutoff-thresholds required the ratio of positive to negative cells as an additional hyper-parameter (**Figure 2C and S1B**). Further, we found that large-scale intensity gradients cause false-positive detections in brighter regions and false-negative detections in dimmer regions of the image, for classifications based on LFI (**Figure 2D**). In contrast, the (0, ∞)-interval of the LFC-distributions aligned with the expected number of positive cells, and the LFC readout accurately represented the spatial distribution of antibody-positive cells, even in cases of large-scale intensity gradients (**Figure S1C**). In addition to spatial dependences within a single experiment, batch effects are common when jointly analyzing multiple experiments. For both LFI and LFC, data-integration was improved by filtering entries of the feature matrix by a fixed minimum threshold. However, low-dimensional UMAP embedding of the filtered LFC showed large overlap between cells from two separate CODEX experiments, while the filtered LFI retained its experimental bias (**Figure 2E**). That finding is in line with our reasoning that due to local normalization, the LFC is inherently less susceptible to experimental batch effects.

In order to investigate the influence of filtering on the subsequent analysis, we varied the minimum LFC- and LFI-filter cutoff and assessed the effect on downstream analysis steps. To assess whether the filtered feature matrix aligns with the expected antibody abundance, we calculated the antibody-wise sparsity as the fraction of zero-entries in each column of the feature matrix. With increasing filtering cutoff, the antibody-wise sparsity increased in a sigmoidal fashion (**Figure 2F**). For each antibody, the optimal filtering cutoff should result in a feature-wise sparsity that reflects antibody abundance. While all antibodies inherently exhibited approximately 50% sparsity for LFI>0, the sparsity profile for LFC > 0 is consistent with variable abundance of different antibodies. To remove cells that are not covered by the antibody panel, we discarded cells where more than 90% of antibodies have no signal. As a result, the fraction of cells retained after pre-processing decreases for both normalization techniques, while the number of Leiden clusters increases due to the increased sparsity (**Figure 2G**). Next, we calculated silhouette scores with respect to experimental-derived label as a measure of remaining batch effect (**Figure 2H**). Throughout filtering thresholds, the LFC feature exhibited better or equal dataset integration, in terms of lower silhouette score, compared to the LFI feature.

While local normalization can mitigate the effect of large-scale intensity gradients, different levels of baseline fluorescence and variable signal-to-noise ratio, it cannot differentiate between antibody and autofluorescence signal. Specifically, we found that pixel-based autofluorescence subtraction often does not achieve complete removal (**Figure 2I**), and high correlation indicates that a cell is contaminated with autofluorescence especially in the case of high-intensity antibody signal (**Figure 2J** and **Figure S2A**). To account for that complication, we constructed a quality-control statistic by computing the Spearman correlation coefficient between each antibody signal and the blank image in the corresponding channel, and taking the median (**Figure S2B**). Subsequently, we were able to selectively reject autofluorescence-contaminated cells or antibody signals from individual cells without discarding entire samples or tissue regions (**Figure 2K**). The individual pre-processing steps showed high variability in the effect on the different antibody signals (**Figure S2C-D**). After applying the complete set of pre-processing steps, we found that ranking antibody signals by the median of the pre-processed distribution aligned with a visual assessment of antibody staining quality (**Figure S3A-B**), where greater separation correlated with higher quality in a higher number of cells.

### Automated hierarchical cell-type annotation

Since strong over-clustering is needed to resolve biological signatures, we developed an automated annotation strategy according to predefined cell-type signatures, which are constructed from a user-defined cell-type hierarchy (**Figure 3A and Figure S4A**). Further, our strategy exploits the fact that the LFC feature is calibrated around zero, to achieve classification as antibody-positive or –negative for each cluster (**Figure 3B-C and Figure S4B)**, and indeed we found that the binary cluster classification aligns well with the embedded feature distribution (**Figure S4C-D**). Nevertheless, deriving cell-type annotations from binary classification risks uncontrolled propagation of classification errors. Thus, in order to derive a certainty measure, clusters are tested against each reference distribution separately, and the resulting p-values are converted into a matrix of minimum posterior probabilities (MPP) that reflect cluster signatures independently of antibody-abundance and staining intensity (**Figure 3D**). Employing a matrix multiplication formalism based on cluster-wise MPPs and cell-type signatures, we arrived at the annotation-score matrix. The score matrix is usually sparse, but clusters with multiple high-scoring annotations are found as the result of closely related compartment definitions or poor staining quality (**Figure 3E**). Finally, cell-type annotation is set to the row-wise maximum of the annotation score matrix for each cluster.

**Figure 3:**
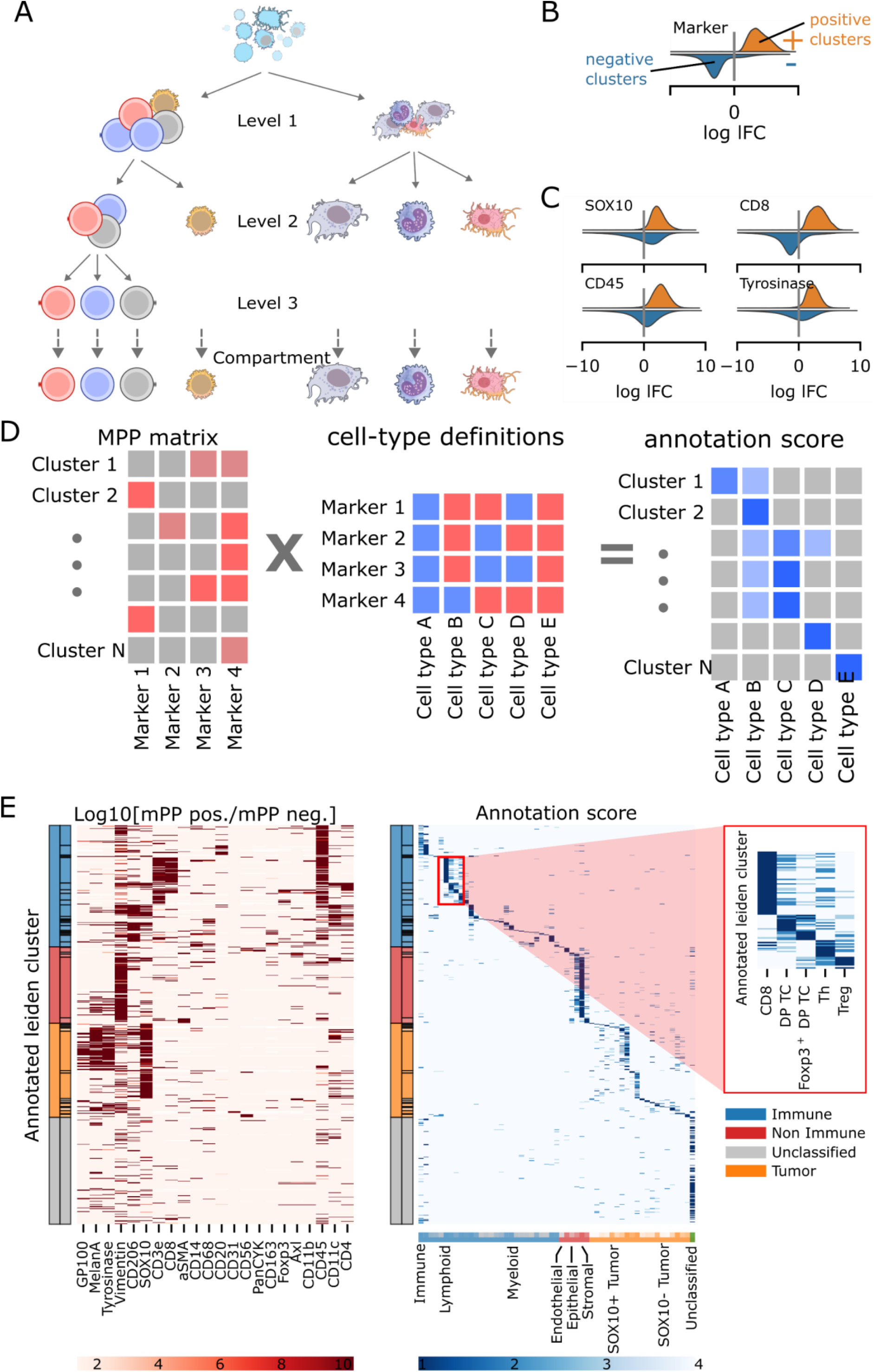
Hierarchical automated cell-type annotation. **(A)** Schematic of hierarchical phenotyping scheme provided as user input. **(B-C)** Schematic and representative examples of positive and negative reference distributions for representative antibody signals used for cell-type annotation, full set shown in Figure S3A. **(D)** Cluster-wise annotation scores are constructed using a matrix-multiplication formalism based on a user defined cell-type hierarchy (‘phenotype definitions’, cf. panel A and S3B) and the minimum posterior probability (mPP) matrix. **(E)** Log-probability ratio of positive vs. negative mPP (left) and combined annotation score (right) of the analyzed melanoma TMA data set. Row-wise rank transformation was applied for visualization of the annotation score matrix. The inset illustrates a set of phenotypic annotations resulting from the work-flow in terms of row-maximal annotation score.

Overall, by means of an image-level normalization scheme applied after cell segmentation, we constructed a single-cell readout that accurately captures the cell-based antibody signals and robustly integrates multiple experiments. Based on that quantity, we constructed an automated annotation approach, thus yielding an end-to-end pipeline down to cellular phenotyping without the need for manual intervention.

### Immune-cell phenotyping of a first-line ICB melanoma cohort

To study the association of *in situ* single-cell features and spatial patterns with response to immunotherapy, we obtained a PhenoCycler dataset with 37 fluorescent channels from two melanoma TMAs comprising 24 patients under first-line ICB therapy (**Figure 4A-B**)(cf. Figure 1D-E). Tissue samples were collected at baseline and subjected to CODEX imaging prior to ICB therapy with anti-PD1, anti-CTLA4 or combination treatment. 14 patients were classified as partial or complete responders (R) and 10 as progressive disease, that is non-responders (NR) to ICB therapy (ORR = 58%), and treatment response substantially improved patient survival (**Figure 4C**). SPARQ-MI’s local feature normalization allowed us to integrate samples from the two separate experiments, which were pooled and jointly analyzed, yielding a data-set of 292 271 annotated cells, on average 12 100 per patient **(Figure 4D, Figure S5A**). We found that the broad tissue architecture is determined by the distribution of myeloid, lymphoid and tumor cells. While we identified regions with predominant occurrence of tumor or immune cells, also the tumor regions show substantial immune-cell infiltration. The experiments comprise skin samples, lymph node or other soft-tissue metastasis, and individual samples in the dataset exhibit a high degree of variability in their microanatomy.

**Figure 4:**
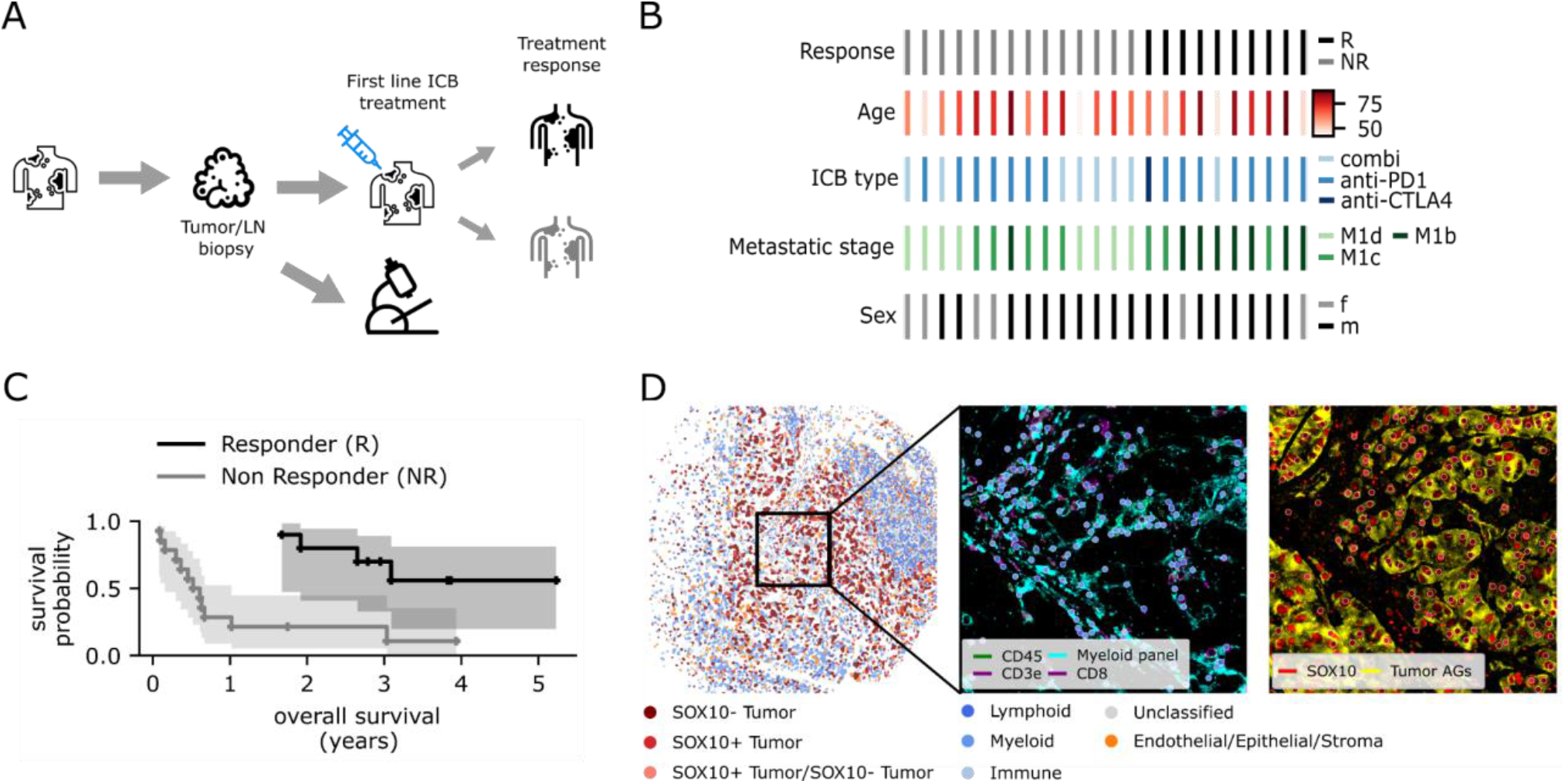
Codex multiplexed imaging of a stage IV melanoma ICB cohort. **(A)** Sample procurement scheme. Primary tumor or lymph node biopsies were collected at baseline from treatment of naïve patients and subjected to CODEX imaging. Patients subsequently received ICB therapy and treatment response was assessed. **(B-C)** Cohort characteristics and Kaplan-Meier estimate of survival probability stratified by ICB response, R: Partial or complete responders; NR: non-responders, that is progressive disease. **(D)** Segmentation mask and cell types for a representative tissue core, and representative cell-type annotation in a region-of-interest at the center of the tissue-core split by immune and tumor cell types.

Our highly automated workflow resulted in quantification of cell-type occurrence, with resolution down to subpopulations such as T cells of type CD8/CD4/Treg, as well as M1/M2 macrophages and several types of tumor cells (**Figure 5A and B**). Moreover, the workflow allowed for in-depth characterization of cell states within those subpopulations, such as cycling (Ki67+) or cytotoxic (GranzymeB+) CD8 T cells (**Figure 5C and D**). While stromal cells accounted only for a small proportion of the TME, myeloid cells, composed in similar proportions of macrophages, dendritic cells and monocytes, outnumbered the lymphoid lineages in the melanoma TME. B-cell contribution to the lymphoid compartment was rare, and myeloid cells are dominated by M2-like macrophages. Most tumor cells expressed the canonical melanoma marker SOX10, and SOX10^-^ tumor cells could still be characterized by expression of different melanoma antigens and constituted a measurable component of the tumor compartment (cf. Figure S4A). The marker Ki-67 indicating cell-cycle progression was detected in tumor and immune cell-types at similar proportion, while GranzymeB and NKG7 expression, indicating cytotoxicity, is restricted to the immune-cell compartment. The immunosuppression-linked marker CD73 (Giraulo *et al*, 2023; Reinhardt *et al*, 2017; Turiello *et al*, 2022) is prominently expressed on non-immune/non-tumor cells, and was detected in immune and tumor lineages as well.

**Figure 5:**
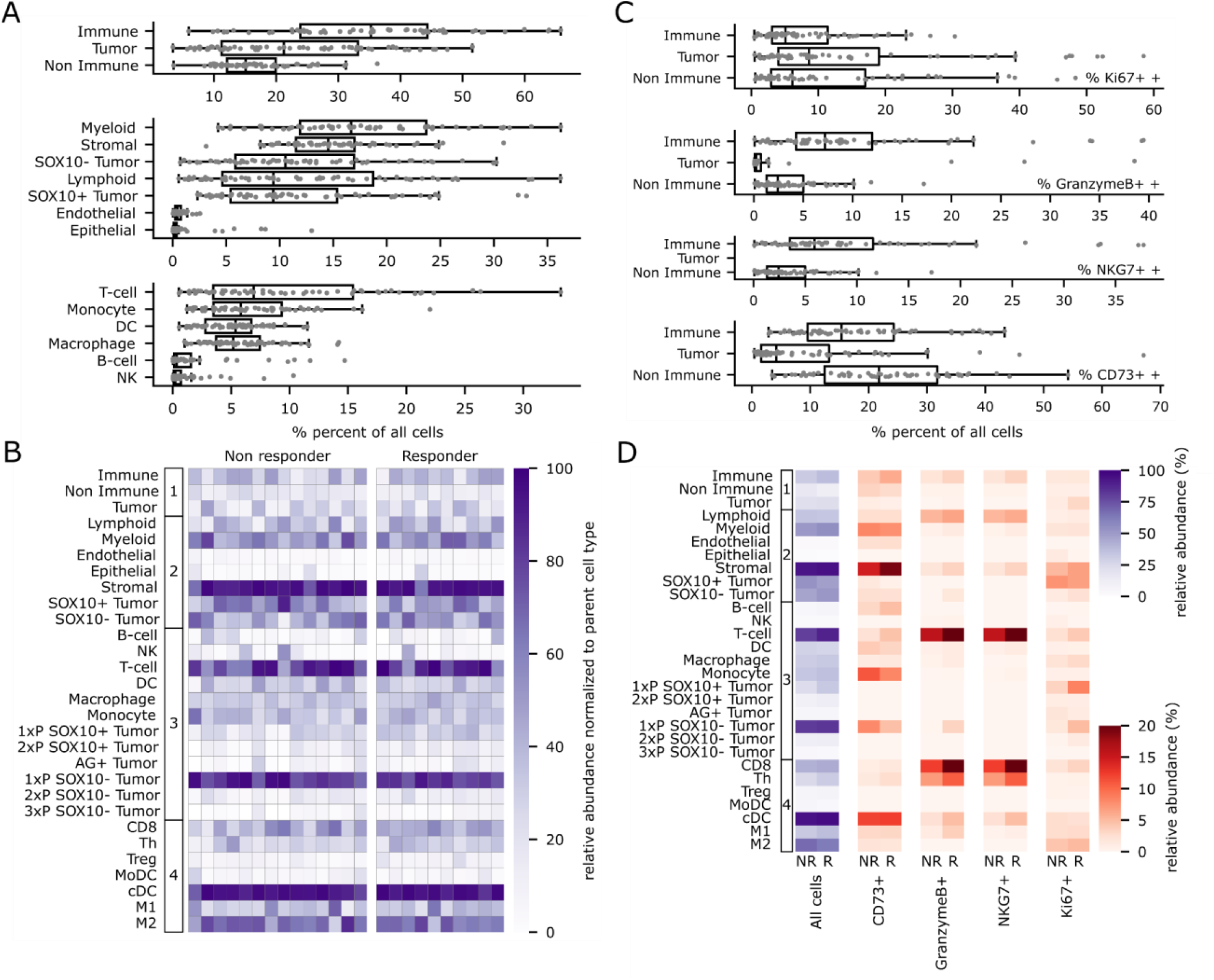
Quantitative analysis of cell types and states. **(A)** Distribution of relative cell-type abundance at annotation level 1 (immune/tumor/non-tumor), 2 and 3. Dots represent TMA cores. (**B)** Patient-level heatmap of relative cell-type abundance aggregated by ICB response. Abundances are normalized to their parent level annotation, for instance the entry for CD8 T cells represents the fraction of that subtype within T cells. (**C)** Occurrence of different cell-state markers aggregated at annotation level 1. Dots represent TMA cores. (**D)** Occurrence of immune-suppressive (CD73), cyto-toxic (GranzymeB, NKG7) and cycling (Ki67) cell-states in all cellular compartments aggregated by ICB response.

### Neighborhood tissue-segmentation allows assessment of immune-cell infiltration

Across the whole dataset, cell-type proportions show a high degree of variability caused by microanatomy, biological variability and TMA composition (cf. Figure 5A and C). To address the effect of tissue architecture as a confounding variable, we developed a modified cellular-neighborhoods approach to segment the TMA cores into immune, tumor and tumor-boundary regions (**Figure 6A**). Based on the resulting segmentation mask, we calculated cell densities and tumor-infiltration depth using the signed distance function of the tumor outline (**Figure 6B**). That tissue-segmentation approach accurately captured the tumor region, allowing separate analysis of tumor and immune regions (**Figure 6C and Figure S5B-C**). The ratio of immune-to-tumor regions varied across TMA cores and did not show significant correlations with ICB response (**Figure 6D-E**).

**Figure 6:**
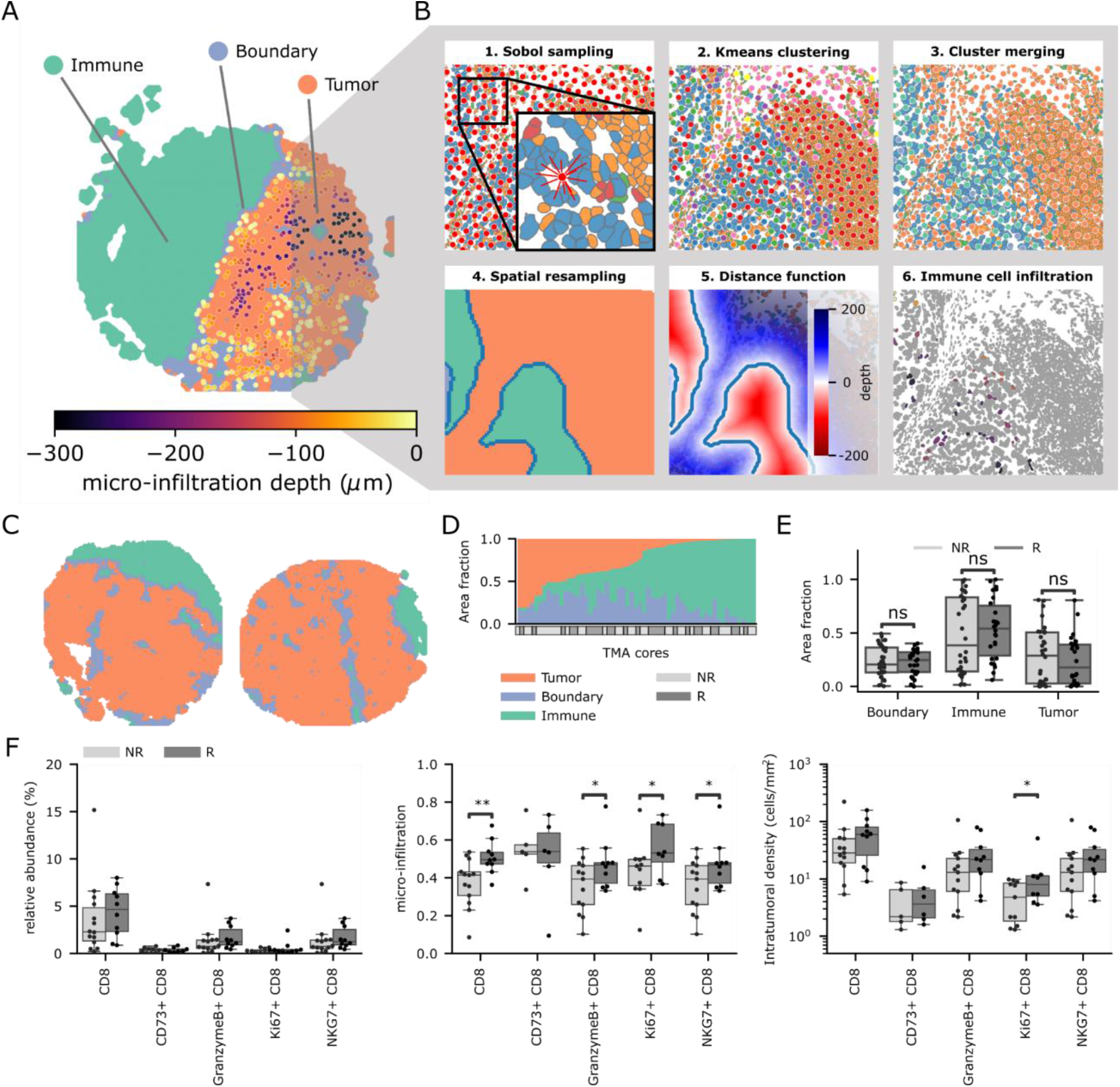
Point-pattern based region segmentation. **(A)** Representative tissue core segmented into immune, inflamed boundary and tumor regions. **(B)** Workflow-schematic of Neighborhood-based region segmentation. **(C)** Representative result of the region segmentation on a subset of tissue cores (full set provided in Figure S4B). **(D-E)** Relative area occupied by different tissue compartments for each patient, sorted by ascending fraction immune compartment area (D) and comparison between different origin tissues (E). **(F)** Micro-anatomically stratified abundance and localization CD8 T cell phenotypes grouped by ICB response. Relative abundance (left), normalized micro-infiltration (middle), density in the intra-tumoral space (right); Dots indicate individual patients and stars indicate significance computed from a Mann-Whitney-U test(* p < 0.05, ** p < 0.01).

Aggregating the infiltration depth and density statistics by the ICB response allowed a refined analysis of the contribution of CD8 T cells to ICB response across patient samples **(Figure 6F)**. Relative abundance of GranzymeB+ and NKG7+ CD8 T cells were significantly increased in responder compared to non-responder samples. Micro-infiltration and intratumoral density showed the same trend, and in addition, all CD8 T cells as well as cycling (Ki67+) CD8 T cells were significantly higher. Interestingly, extratumoral density (**Figure S5D)** did not show any significant differences.

Overall, using SPARQ-MI, we analysed a complex tumor tissue data-set consisting of two separately obtained tissue microarrays. Single-cell feature analysis yielded an overview of aggregated cell-states across patients, which was substantially refined after accounting for different tissue types by employing a customized tissue-segmentation approach.

## Discussion

We constructed an open source end-to-end framework for the processing and analysis of multiplexed imaging data from complex clinical samples, which unifies the processing steps for multiplexed images in a single implementation. As a central element downstream of cell-segmentation, we propose LFC feature normalization at the image level, to achieve calibrated single-cell readouts that are robust to spatial inhomogeneities in the signal distribution. The LFC feature showed a more gradual decrease in cell numbers compared to MFI, as well as a smoother increase in the number of Leiden clusters, allowing for improved control over the pre-processing performance. Further, this approach achieved high experimental integration and allowed for direct classification of marker-positive cells. SPARQ-MI is available as a python library and all dependencies are under open-source licenses.

Building on the calibrated LFC feature, automated phenotyping allowed for high clustering resolution resulting in deep phenotypes without manual intervention. Alternatively, supervised deep-learning methods were proposed for end-to-end processing of multiplexed imaging data (Brbić *et al*, 2022; Sultan *et al*, 2025), but the need for supplying training data makes supervised methods costly as a standard protocol for cellular phenotyping. Unsupervised methods (Geuenich *et al*, 2021; Zhang *et al*, 2022; Amitay *et al*, 2023) treat phenotyping as a statistical problem, enabling the automatic annotation of cell types without manual intervention. Notably, the automated nature of our annotation approach allows for systematic optimization of parameters even for processing steps upstream of cluster annotation, as it treats the entire phenotyping scheme as a compound hypothesis rather than assessing individual cell types in isolation.

For spatial analysis, using a neighborhood-based region segmentation approach (Patrick *et al*, 2023; Chen *et al*, 2020) enabled the weakly supervised detection of micro-anatomical tissue compartments. That approach parallels recent workflows of tissue segmentation for spatial transcriptomics splitting the tissue section into distinct regions (Yuan *et al*, 2024), which however are not directly applicable to lower-dimensional data types such as spatial proteomics. Our neighborhood-based implementation requires only annotated cell types as input, and allows for convenient calculation of readouts such as cell density and tumor infiltration depth.

In our cohort of stage IV melanoma patients under first-line ICB treatment, statistical modeling based on spatial features and high-level TME states proved a robust approach to analyze immune-cell phenotypes in a clinical context. By accounting for the micro-anatomy of the TME, we were able to recover the beneficial predictive effect of cytotoxic (GranzymeB+) CD8 T cells on ICB therapy response from a heterogenous dataset. Furthermore, our analysis reveals an association of the proliferation marker Ki65+ on T cells in the tumor with improved response to immunotherapy, in agreement with previous studies in the context of cancer and also autoimmune disease(Edwards *et al*, 2025; Huang *et al*, 2024; Wang *et al*, 2023; Horn *et al*, 2025).

Given the numerous signaling and regulatory interdependencies between immune-cell subtypes, a sophisticated understanding of the TME requires the simultaneous analysis of many cell types in their native spatial context(De Visser & Joyce, 2023; Lu *et al*, 2025; Balaban & Cohen, 2025). For instance, in strongly infiltrated tumors, pro-inflammatory immune cells create a cytokine milieu that promotes the activity of anti-tumor effector cells, and suppressive immune-cell phenotypes can create a local environment that inhibits tumor infiltration. Recent work has proposed systematic statistical analysis and machine-learning techniques as well as mechanistic mathematical modeling workflows, to investigate the tightly regulated spatiotemporal dynamics of immune-cell populations (Steinheuer *et al*, 2025; Lee *et al*, 2024; Van Santvoort *et al*, 2025; Agrawal *et al*, 2024). Such methods require quantitative spatial information as input, and hence a streamlined and efficient computational end-to-end workflow for multiplexed image analysis.

Overall, SPARQ-MI presents a scalable, integrated framework for analysis of multiplexed histology data from complex clinical tumor samples, with potential for statistical inference of single-cell and spatial features and for deriving a quantitative understanding of immune-cell organization in the environment of tumor biopsies.

## Methods

### Patient Cohort

Two tissue microarrays (TMA) comprising metastatic samples from 24 patients with advanced melanoma were used. The TMAs have been previously described (Braun *et al*, 2020)(“Bonn cohort”). Specimen were obtained from the Department of Dermatology and Allergy of the University of Bonn (Germany) and preserved as formalin-fixed paraffin-embedded (FFPE) tissues. The TMAs were prepared including three tissue cores for each patient. Sections of 4 µm thickness were obtained and mounted on poli-lysin coated microscope coverslips prior to staining procedures. All samples were collected before initiation with anti-PD-1 monotherapy (nivolumab or pembrolizumab), anti-CTLA-4 (ipilimumab), or anti-PD-1 in combination with anti-CTLA4. All patients signed an informed consent, in accordance with participating hospitals/research institute Human Research Ethics Committee procedures and guidelines and conforming to the Declaration of Helsinki.

### Multiplexed immunofluorescence imaging

Multiplexed immunofluorescence experiments were performed as we previously described(Giraulo *et al*, 2023). Image acquisition in sequential cycles was performed using the Phenocycler® instrument (Akoya Biosciences). For each cycle the instrument enables the acquisition of up to three oligonucleotide-conjugated antibodies together with the nuclear counterstain DAPI(Black *et al*, 2021). The DNA oligonucleotides were purchased from <u>Biomers</u>. The antibodies were conjugated to the corresponding DNA oligonucleotides, following the protocol already published by others(Black *et al*, 2021), that we slightly adapted. Images were acquired using a Zeiss Axio Observer 7 inverted microscope, equipped with Colibri 7 as the LED Light source, the Plan-Apochromat 20X/0,8 M27 (a=0,55 mm) as objective and the Prime BSI PCIe camera. The following LED intensity and exposure time settings were used: DAPI 40% intensity, 20 milliseconds (ms); Cy3 65% intensity, 400 ms Cy5 50% intensity, 500 ms; Cy7 50% intensity, 500 ms. Once acquired, the images were converted to TIF files by using Codex Instrument Manager® (Akoya Biosciences). The human cholangiocarcinoma dataset has been previously published(Branchi *et al*, 2024). Multiplexed immunofluorescence of the murine spleen was performed following the protocol described in another study(Van Der Voort *et al*, 2026).

### Image pre-processing and segmentation

The image pre-processing pipeline takes raw image data from the CODEX Uploader software and outputs stitched best-focus multichannel images. Optionally final images can be obtained as ome.tiff. Images are processed using deconvolution(Czech *et al*, 2019), focus merged using the extended depth of field approach(Forster *et al*, 2004). Tiles are registered and cycles are aligned using Ashlar(Muhlich *et al*, 2022). We extended the original background subtraction scheme(Goltsev *et al*, 2018) by rescaling the blank image for each cycle based on a correction calculated from high auto-fluorescence regions. For a single channel the correction is calculated as follows. First, we calculated an autofluorescence image *B* as the maximum projection of blank images in the first and last cycle. Subsequently, a set of probe locations *P* is randomly drawn from the upper 10 percent of the autofluorescence-channel intensity distribution. We assume that the fluorescence signal at the probe locations will be dominated by background signal throughout the experiment. This allows us to calculate a scaling factor for each cycle *i* as the ratio between fluorescence signal and autofluorescence at the probe locations

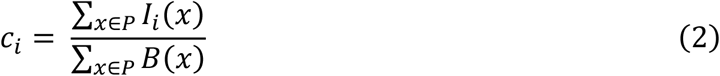

The autofluorescence image is rescaled for each channel prior to subtraction

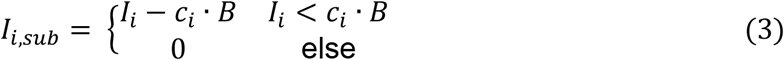

Cell segmentation was performed using the Mesmer(Bannon *et al*, 2021) model for murine spleen tissue and Cellpose(Pachitariu & Stringer, 2022) for clinical tumor samples. The baseline Cellpose model was fine-tuned using dataset specific training data.

### Local feature normalization

Single-cell fluorescence intensity features were normalized to the antibody-specific local noise level as follows. For each antibody, we compute (i) a band-pass filtered foreground *I*_*f*_ and (ii) a high-pass filtered image background image *I*_*b*_, where the band-pass filter is chosen in an way to capture subcellular structure and the high-pass filter cutoff is chosen in line with the optical resolution of the imaging platform. A normalized single cell feature is computed as the local log2 fold-change (LFC, **Equation** 1) between the average band-pass filtered foreground and high-pass filtered background intensities The foreground distribution *F*_*i*_ is given by the pixel values belonging to each cell footprint *F*_*i*_ = {*I*_*f*_(*k*)| *k* ∈ *M*_*i*_}, where *M*_*i*_ is the set of pixels belonging to the segmentation mask of the *i*-th cell. The background distribution *B*_*i*_ is sampled from an exponentially weighted environment of each cell

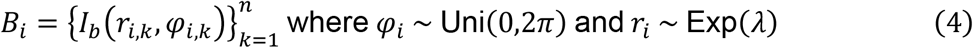

where *r* and *φ* are coordinates in polar reference frame centred around each cell and the number of samples points *n* as well as the size parameter *λ* are hyper-parameters. Additionally, one-sided p-values for the LFC were calculated using a Mann-Whitney-U test under the null hypothesis that *F*_*i*_ > *B*_*i*_.

### LFC vs. LFI parameter scan and batch-effect quantification

Minimum LFI and LFC thresholds in the single cell pipeline described below were varied from -5 to 5, with the filters imposed by U-statistic, p-value and QC-statistic disabled. Antibody-wise feature sparsity was calculated as the fraction of cells with a non-zero value. The number of clusters was calculated by applying Leiden clustering to the resulting dataset for both LFI and LFC separately. The silhouette score with respect to the exp ID label in high dimensional and UMAP space was used as a measure of remaining batch effect(Büttner *et al*, 2019).

### Single cell autofluorescence quality control statistic

The autofluorescence quality control statistic is calculated from the single-cell correlation coefficient between the pixel intensities of the antibody signal and the corresponding blank image. (i) We compute the spearman correlation coefficient between the pixel values for a single antibody marker and the pixel intensities of the blank cycle image in the corresponding fluorescence channel. This calculation is repeated for each cell. The resulting correlation matrix has the same structure as the single-cell feature matrix. (ii) The median correlation coefficient across all antibodies is computed per cell. The resulting statistic is used to filter out entire cells in the pre-processing based on a fixed cutoff. Similarly, individual entries of the feature matrix can be filtered based on the corresponding correlation coefficient.

### Single cell pre-processing, embedding and clustering

The segmented objects enter into the single cell processing as a feature matrix where each column corresponds to a feature and each row represents an object. The feature matrix is filtered by either removing entire objects or setting individual entries to zero. Cell and features are filtered by maximum allowed QC-statistic to remove the effect of residual autofluorescence. The p-values calculated during feature extraction are used to remove noisy detections by filtering features based on minimum log p-value. To increase the sparsity of the dataset for subsequent clustering, the man-Whitney-U statistic and log_2_(LFC) are used to remove low intensity detections. Outliers resulting from very bright fluorescent debris is removed by imposing a maximum log_2_(LFC) threshold. Subsequently, cells with fewer than n nonzero features are removed. Fluorescence intensity features are normalized using z-normalization and logarithmized. Morphological features are z-normalized. After pre-processing the filtered feature matrix is clustered using the Leiden graph partitioning algorithm and UMAP projections are generated based on a joint k-nearest neighbor graph as implemented in Scanpy(Wolf *et al*, 2018).

### Phenotyping scheme and automated cluster annotation

The phenotyping scheme is expressed as a tree where the nodes are cell types or phenotypes and the edges represent cell type - subtype relationships. Each node may have a list of lineage defining positive, negative and excluded markers, that are represented in the phenotyping matrix by 1,-1 and 0 respectively. To create concrete signatures, negative markers are inherited to all children of a node, while positive markers are propagated upwards from children to parent nodes. The individual signatures form the columns of the phenotype matrix

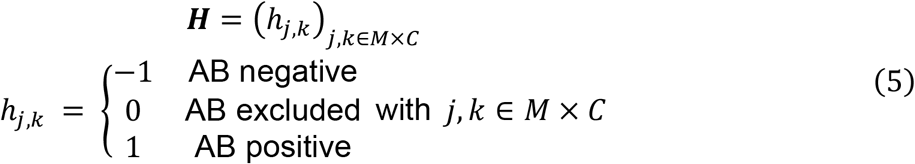

where *C* is the set of all phenotypes and *M* is the set of all markers. Additionally, each node is associated with an annotation level that is given by the distance to the root node. The hierarchical structure is used to conveniently create individual cell type signatures but does not have an impact on the phenotype scoring, however the node level will be used to resolve ties.

Leiden clusters are categorized as positive or negative based on their 10%-percentile value for each antibody. To generate a confidence scores of the +/-assignment for each cluster, we estimate assignment probabilities as follows. First, reference distribution for each antibody is constructed from the union of all cells in positive and negatively classified cluster, respectively

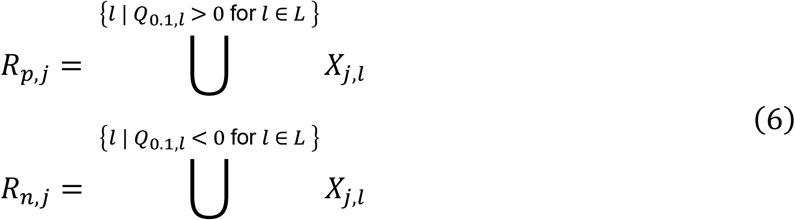

Where *L* is the set of all Leiden clusters and *X*_*j,l*_ represents the feature distribution of a given cluster *l* and antibody *j*. Subsequently each cluster is tested separately against the positive and negative reference distribution using a non-parametric statistical test by testing the hypothesis that *R*_*p,j*_ = *X*_*j,l*_ and *R*_*n,j*_ = *X*_*j,l*_.

The resulting p-values are converted into minimum posterior probabilities using the SBB approach(Held, 2010; Sellke *et al*, 2001)

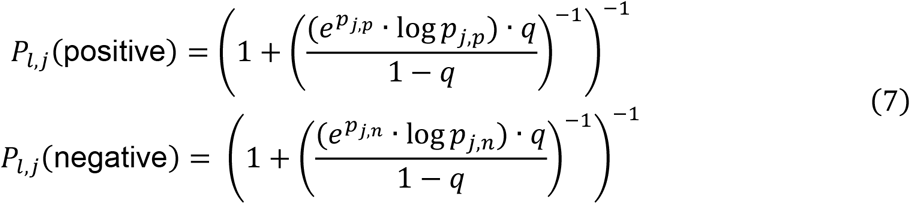

where *p*_*i,p*_ is the p-value from testing against *R*_*p,i*_. This yields a probability matrix where each entry is associated with a cluster *l*, antibody *j* and either positive or negative assignment probability 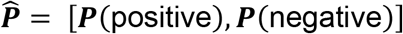. The component matrices are given by

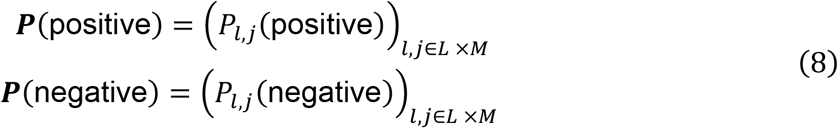

Similarly, the complete phenotype matrix is constructed by concatenating ***H*** and its element-wise additive inverse 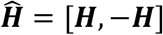. To select the most likely annotation for each cluster a score matrix is computed as the matrix product between the probability matrix and phenotype matrix.

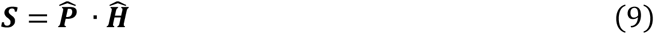

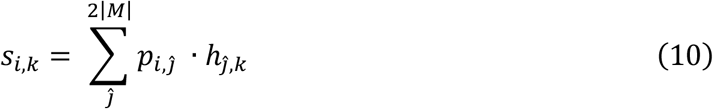

The final annotation is selected as the row-wise maximum of the score matrix. Ties are resolved by choosing the annotation with the smallest number of positive markers, the largest number of negative markers and the smallest annotation level. This ensures that cluster annotations are parsimonious but still allows clusters to receive an annotation even if they do not fit any of the most fine-grained cell types.

### Point pattern region segmentation

Tissue region segmentation were generated using modified cellular neighborhoods (Schürch *et al*, 2020) approach. In brief, feature vectors are constructed from a k-nearest-neighbor network between cell centroids and randomly sampled points chosen through Sobol sampling. We then compute the relative abundance of each cell annotations in the k-neighborhood of each sampling point. The feature vectors are subsequently clustered using K-means. By concatenating feature vectors for different values of k prior to clustering, multiple length scales may be considered. To generate annotations for immune, boundary and tumor regions clusters were merged and annotated based on the abundance of Tumor cells. The resulting labeled point cloud was resampled onto a grid and rescaled to match the dimensions of the underlying cell coordinate system.

### Tumor infiltration statistics

To calculate the infiltration depth of immune cells into the tumor region, signed distance functions (SDF) are calculated, based on the boundary pixels between the immune region and the combination of the tumor and tumor boundary regions, using the *scikit-fmm* library. Single-cell infiltration values are calculated by averaging the SDF corresponding to the tumor region over the pixels occupied by each cell. To make the quantitative relationship between intratumoral and extratumoral cell densities more accessible to downstream models, we defined the infiltration 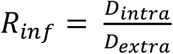 and exclusion 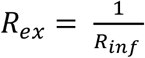 ratios as additional readouts. Normalized infiltration depth is calculated by applying quantile normalization. To analyze tumor infiltration with respect to treatment response and patient survival, relative cell abundance, density and normalized infiltration depth statistics are averaged for each donor. The resulting feature matrix contains normalized infiltration depth, infiltration/exclusion ratios, intra-tumoral and extra-tumor cell infiltration densities as well as fractional abundance for all cells of a given cell type. Additionally, the same features are computed for Ki67^+^, CD73^+^, Granzyme B^+^ and NKG7^+^ single positive subsets. For univariate statistics, missing values were dropped.

