## Supplemental Figures for "SPARQ-MI leverages end-to-end spatial single-cell analysis of the tumor microenvironment"

### Supplementary Figures

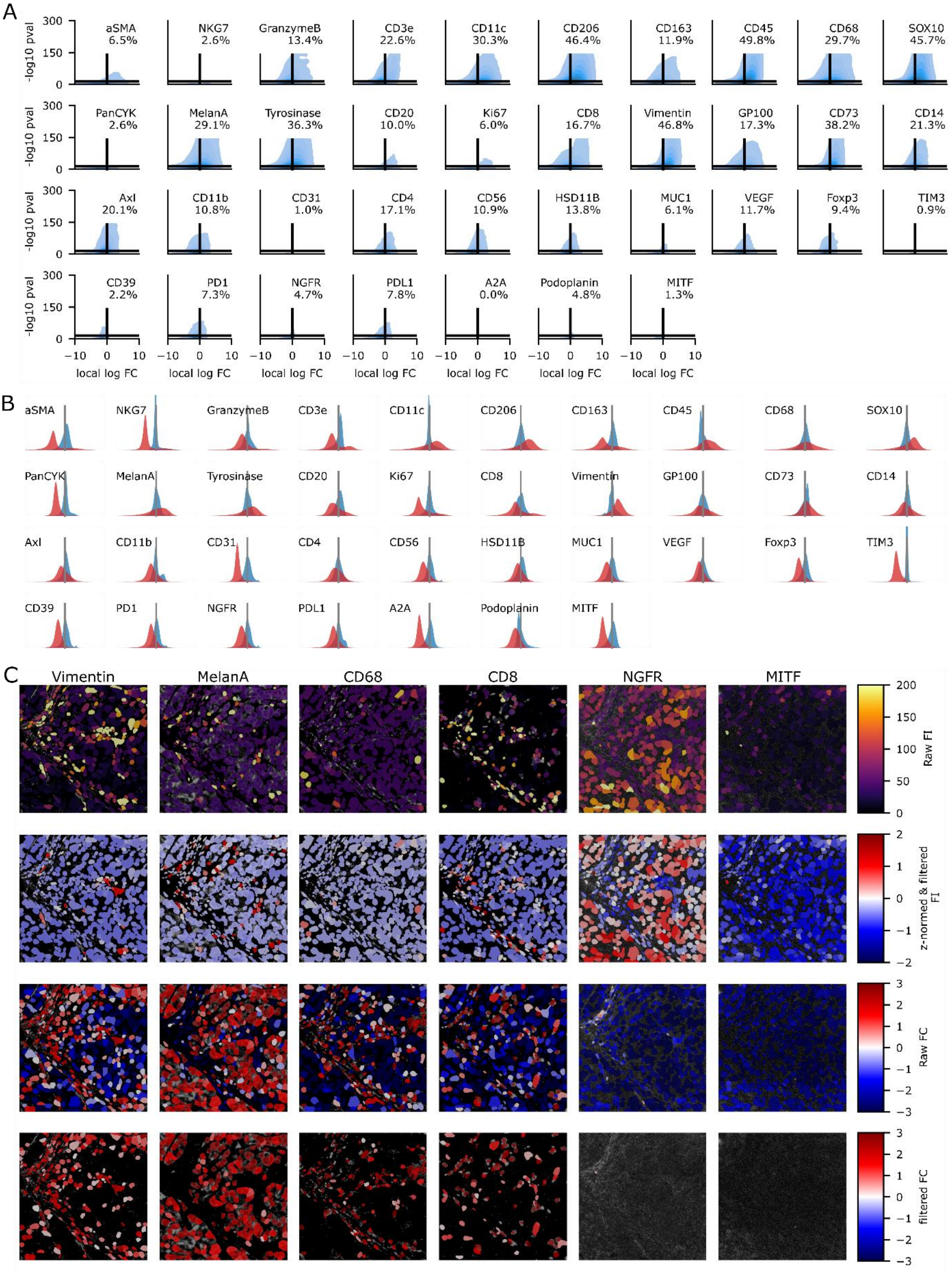

#### Figure S1: Locally normalized feature extraction.

**A:** Kernel density estimates of the LFC features and their associated p-values for each antibody in the melanoma TMA data-set. Percentages indicate the fraction of cells that exceed both LFC and p-value cutoffs in the pre-processing.

**B:** Extended Figure 2C. LFI (blue) and LFC (red) distributions for all antibodies.

**C:** LFI and LFC single cell features before and after pre-processing visualized in a representative tissue region for Vimentin (stroma), MelanA (tumor), CD68 (macrophage), CD8(T cell), NGFR (tumor, low quality staining) and MTIF (tumor, low quality staining) antibodies.

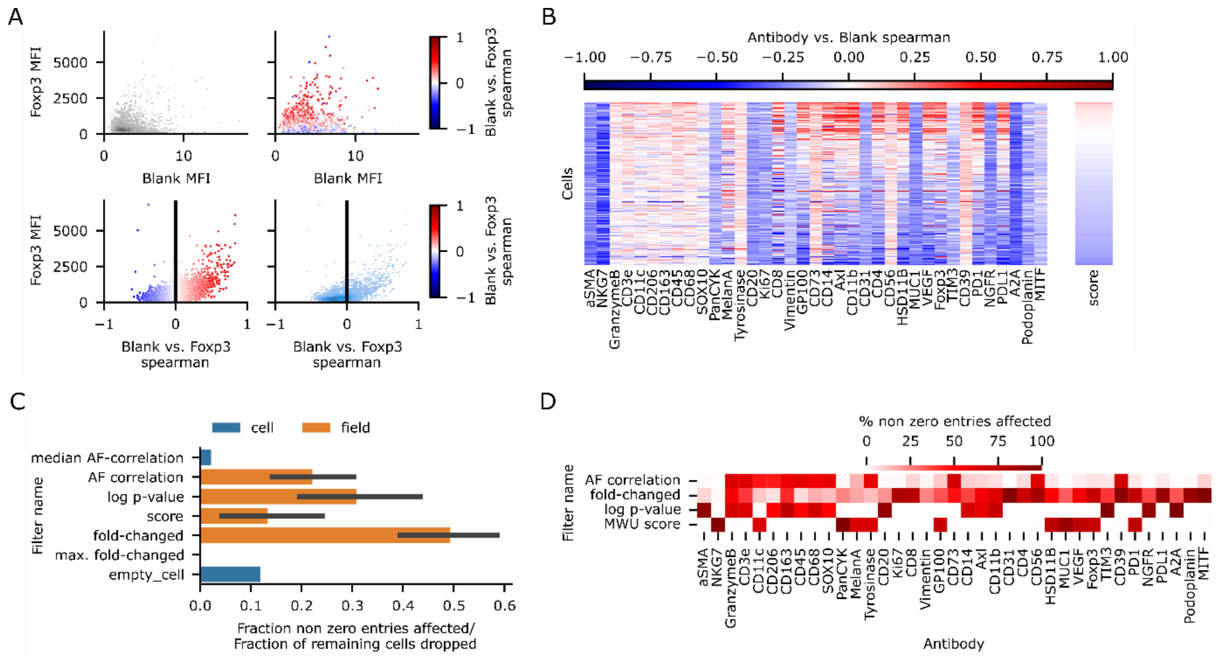


#### Figure S2: Autofluorescence quality control and single cell pre-processing

**A:** Connection between FoxP3 MFI and correlation statistic for a representative tissue core in the melanoma TMA data-set. Top: Correlation between FoxP3 and Blank MFI single cell readout as density (left) and colored by correlation statistic (right). Bottom: Relationship between correlation statistic and MFI as scatterplot (left) and density estimate (right).

**B:** Matrix of blank vs antibody correlation statistic for each cell and antibody. Cells were ranked by median correlation across all antibodies.

**C:** Effect of the individual pre-processing steps. Cell wise filters (blue) discard an entire cell, while element-wise filters (orange) reject individual entries in the feature matrix. Errorbars indicate standard deviation across antibodies.

**D:** Effect of element-wise pre-processing filters split between individual antibodies.

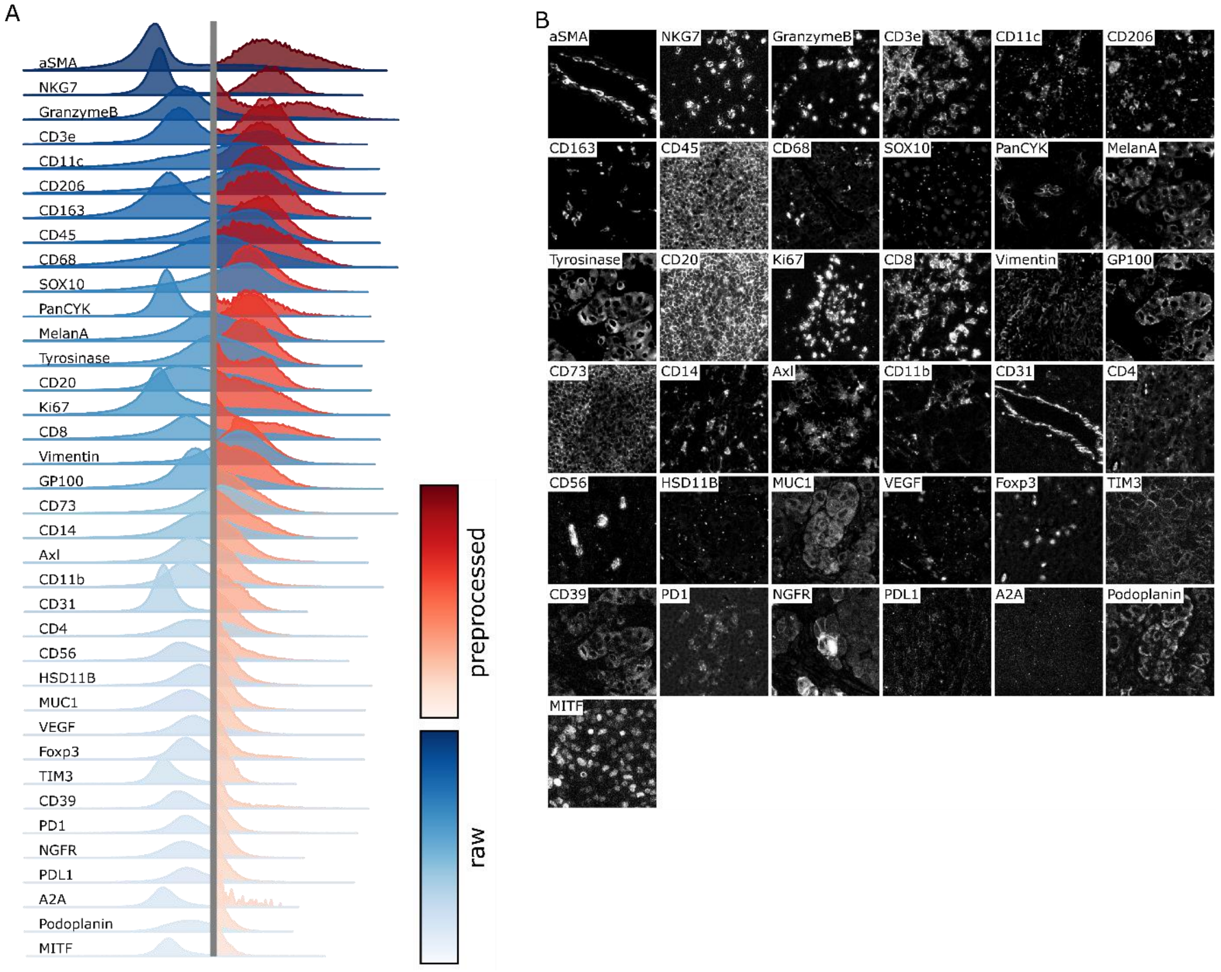


#### Figure S3: Signal distributions and fluorescence images after preprocessing

**A:** Ridge-plot of raw (blue) and pre-processed (red) LFC distributions for each antibody. The grey line indicates LFC=0. Antibodies were ordered by the median LFC of the pre-processed distribution.

**B:** Representative fluorescence images for all antibodies, sorted according to panel S2F. Fluorescence intensity values were rescaled separately for each antibody to the 0% to 99.9% percentile range.

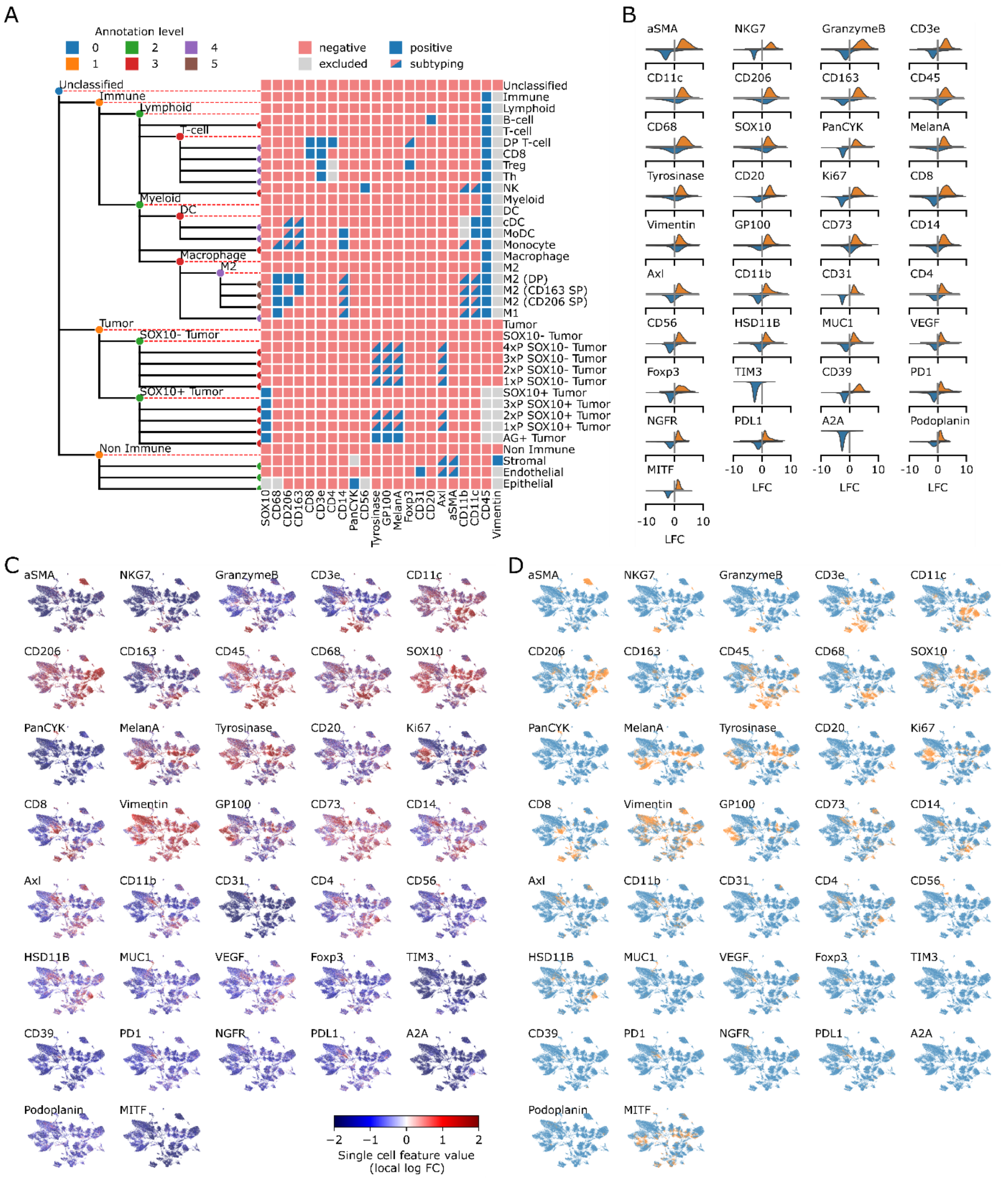


#### Figure S4: Automated cell type annotation

**A:** Truncated cell type matrix linking cell types (rows) to their expected antibody signature (columns). Dashed red lines indicate annotations that are derived from an intermediate branch of the annotation hierarchy. For some cell types, combinations of a marker subset are used to systematically create an ensemble of subtypes.

**B:** Extended version of Figure 3C.

**C:** Raw LFC values for each antibody visualized on the UMAP embedded single cell data.

**D:** Binary pos/neg classification visualized on the UMAP embedded single cell data.

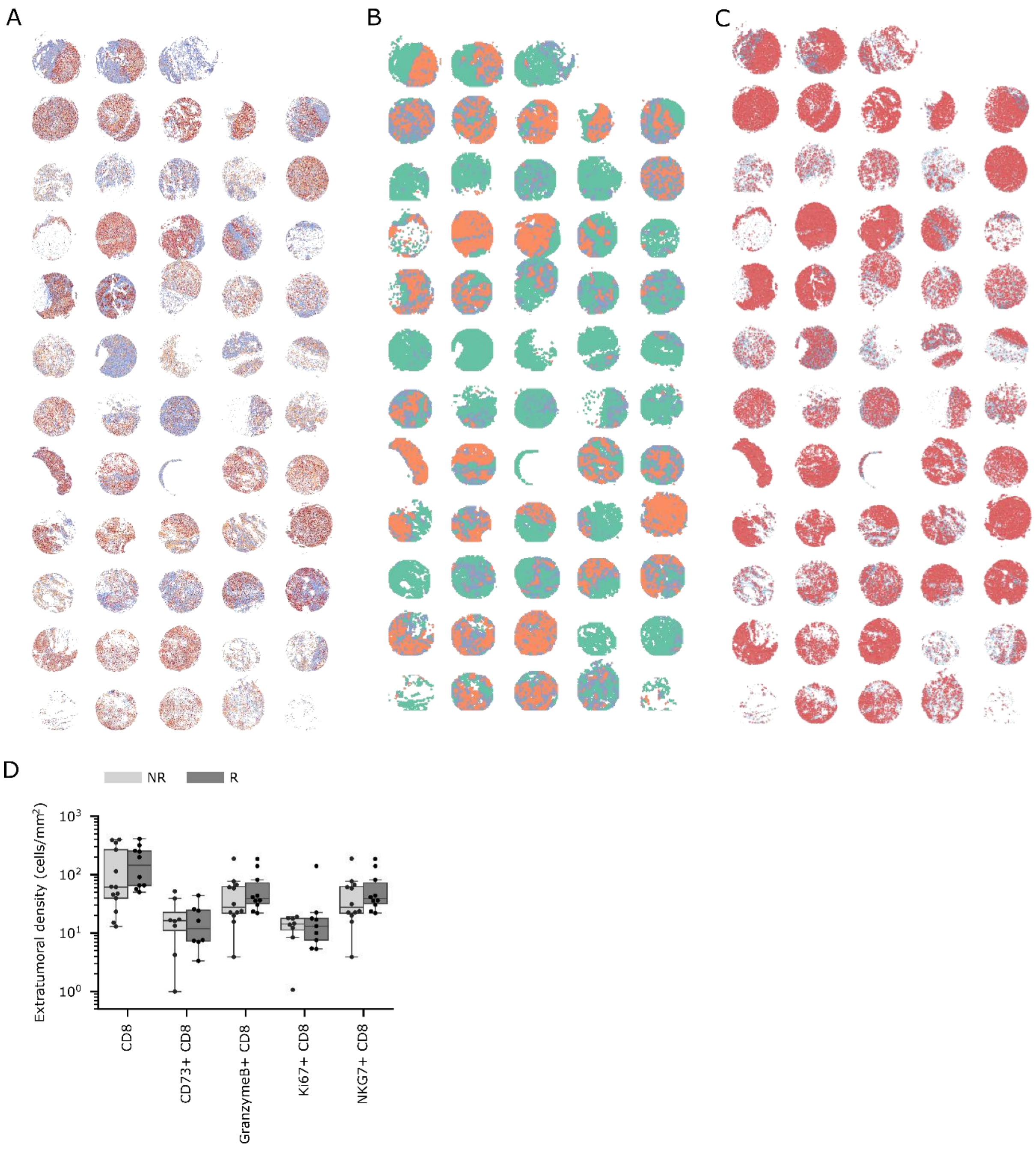


**Figure S5: Visualization and extended analysis of TMA cores**

**A:** Overview of cell-type annotation for all tissue cores, cf. Figure 4D.

**B-C:** Melanoma TMA region segmentation (B, cf. Figure 6C) and corresponding distribution of tumor cells (C, red).

**D:** Density of immune cells in the extra-tumoral space, cf. Figure 6F. Dots represent patient averages.
